# The landscape of cancer rewired GPCR signaling axes

**DOI:** 10.1101/2023.03.13.532291

**Authors:** Chakit Arora, Marin Matic, Pierluigi DiChiaro, Natalia De Oliveira Rosa, Francesco Carli, Lauren Clubb, Lorenzo Amir Nemati Fard, Giorgos Kargas, Giuseppe Diaferia, Ranka Vukotic, Luana Licata, Guanming Wu, Gioacchino Natoli, J. Silvio Gutkind, Francesco Raimondi

## Abstract

We explored the dysregulation of GPCR ligand signaling systems in cancer transcriptomics datasets to uncover new therapeutics opportunities in oncology. We derived an interaction network of receptors with ligands and their biosynthetic enzymes, which revealed that multiple GPCRs are differentially regulated together with their upstream partners across cancer subtypes. We showed that biosynthetic pathway enrichment from enzyme expression recapitulated pathway activity signatures from metabolomics datasets, providing valuable surrogate information for GPCRs responding to organic ligands. We found that several GPCRs signaling components were significantly associated with patient survival in a cancer type-specific fashion. The expression of both receptor-ligand (or enzymes) partners improved patient stratification, suggesting a synergistic role for the activation of GPCR networks in modulating cancer phenotypes. Remarkably, we identified many such axes across several cancer molecular subtypes, including many pairs involving receptor- biosynthetic enzymes for neurotransmitters. We found that GPCRs from these actionable axes, including e.g., muscarinic, adenosine, 5-hydroxytryptamine and chemokine receptors, are the targets of multiple drugs displaying anti-growth effects in large-scale, cancer cell drug screens. We have made the results generated in this study freely available through a webapp (gpcrcanceraxes.bioinfolab.sns.it).

**Significance:** Comprehensive analysis of GPCR extracellular network in cancer transcriptomics datasets reveals signaling axes associated to patient survival, whose targeting is associated with growth inhibition in cancer cell lines drug sensitivity assays.

## Introduction

G-protein coupled receptors (GPCRs) are the largest family of transmembrane proteins which transduce a wide number of physical and chemical stimuli into the cell by coupling to intracellular heterotrimeric G-proteins, thereby controlling multiple downstream signaling pathways and transcriptional programs^1–3^. The GPCRs repertoire expanded and diversified to meet the increasing need for inter-cellular communication in evolving multicellular organisms^4,5^. Systematic efforts are being undertaken to illuminate the complex circuitry of GPCR signaling both at the intracellular ^5–11^ as well as the extracellular ligand level^12^.

Cancer genomics provides the possibility to systematically shed light on cancer somatic alterations affecting GPCRs and their interacting partners ^13–17^. Indeed, a landscape of GPCR-centric autocrine, paracrine, juxtacrine or endocrine signaling networks is emerging as a critical component for the interaction of cancer cells with their tumor micro-environment (TME), ultimately leading to cancer progression and therapy resistance^15,16^. For instance, chemokines, such as *CXCL1*, *CXCL2*, and *CXCL8,* bind and activate their receptors, primarily *CXCR1* and *CXCR2* on immune suppressive myeloid derived cells thereby disabling anti-tumoral immune surveillance mechanisms^15^. Several GPCRs and their associated components mediating inflammation have also been connected to immune evasive TME, such as prostaglandins (in particular *PGE2*), their biosynthesizing enzymes (*PTGS1* and *PTGS2*) and cognate prostanoid receptors (e.g., *PTGER2* and *PTGER4*). Ectonucleotidases (such as *NT5E* or *ENTPD1*) and adenosine receptors are also well known to induce an immunosuppressive tumor microenvironment^15^. Recent studies have also shown the emergent role of innervation in promoting tumor growth (reviewed in^18^). Notably, it has emerged a cross-talk between the neurotrophin pathway, particularly the *NGF-NTRK1* axes, and GPCR signaling mediated by either (nora)adrenaline-*ADRB2* in pancreatic cancer^19^, or acetylcholine-*CHRM1 and CHRM3* in gastric cancer^20^. The neuron-cancer cell communication can also be established via additional routes, such as the one of 5-hydroxytryptamine, which is produced by enteric serotonergic neurons, and activates cognate receptors expressed on colorectal cancer stem cell, which undergo self-renewal and tumorigenesis^21^. However, defining these immune evasive and tumor promoting networks has been elusive.

In this regard, single cell and spatial sequencing technologies are allowing us to study mechanisms of cell-cell communications at an unprecedented resolution^22^. In the most comprehensive resource to date used to infer ligand-receptor pairs mediating cell-cell communication^23^, GPCRs-ligand interactions are surprisingly depleted, representing roughly less than 5% of the total curated interactions, despite being the most abundant class of transmembrane proteins involved in signal transmission. This is likely due to the fact that many GPCRs bind to small organic molecules, including neurotransmitters and multiple metabolites, which don’t represent PPIs detected by most cell-cell communication algorithm, being impossible to be detected via transcriptomics techniques.

In the present study, we considered an extended network of signaling axes formed by GPCRs with interacting ligands, as well as biosynthetic enzyme pathways, to systematically explore their aberrant regulation in cancer transcriptomics datasets and investigate their prognostic value for personalized medicine treatments.

## Results

### GPCRs extracellular signaling network

We first generated a comprehensive catalog of upstream and downstream molecular interactions involving GPCRs (Figure1a-c; see Methods). Briefly, we retrieved known GPCR ligands from IUPHAR/Guide to Pharmacology database (GtoPDB), which we mapped to individual ligand-synthesizing enzymes and to known biosynthetic pathways, as well as to pathways mediating signal transduction (Methods; Supplementary Table 1,2). This yielded a total of 236 ligands, 309 biosynthetic pathways and 82 enzymes associated instances (Figure 1a-c). We found that (86%) of ligands and (99%) of enzymes were already annotated within biological pathway knowledgebases (i.e., Reactome; Figure 1b). A total of 211 (53%) non-olfactory receptors have known ligands in GtoPdb, 95 of which exclusively bind organic ligands, 90 peptides and 26 both ligand types (Figure 1b). While the vast majority of peptide ligands are annotated to participate in signaling pathways, the organic ligands are instead extensively annotated as part of metabolic processes (Figure 1b; Supplementary Table 1).

**Figure 1.**
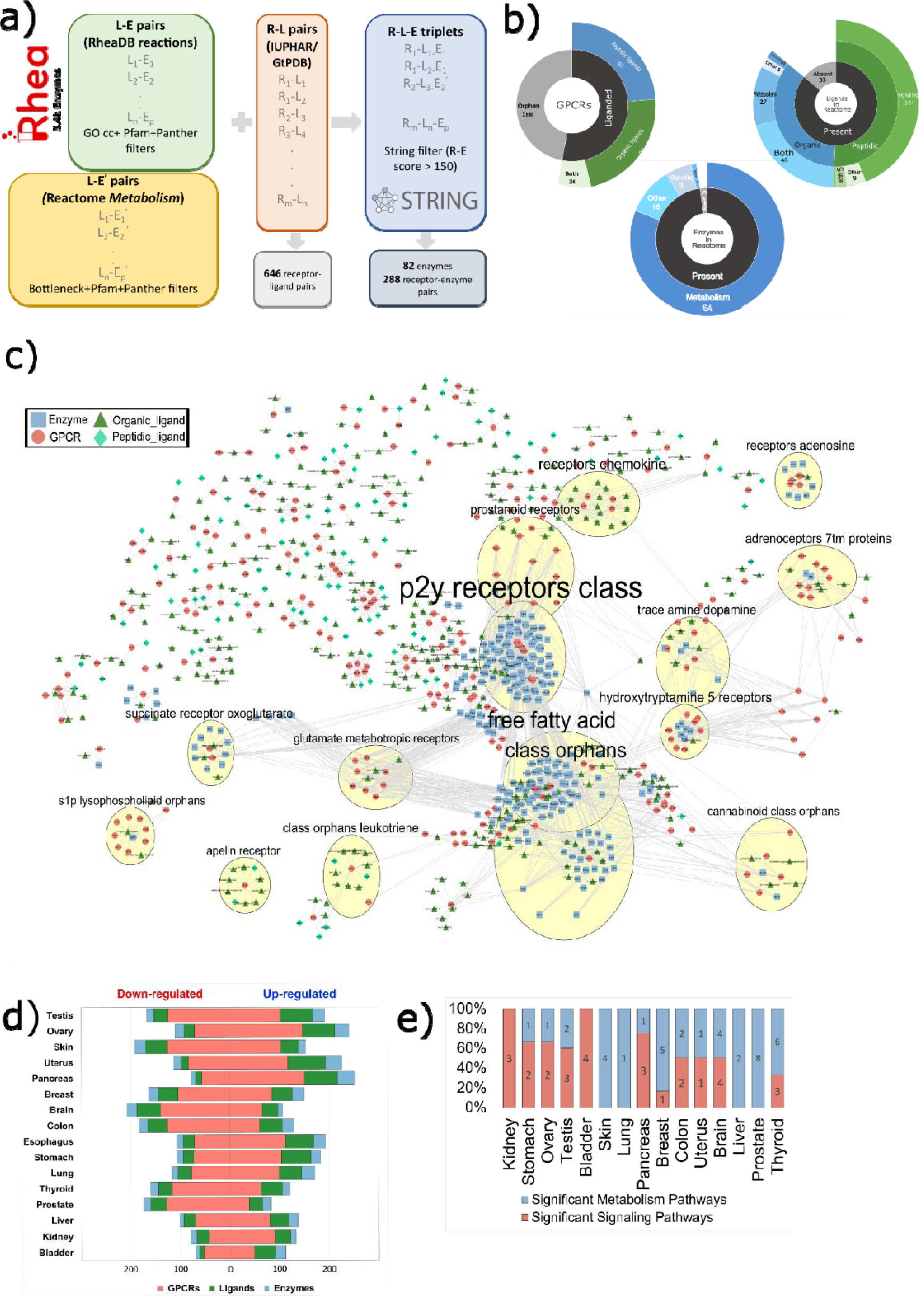
GPCR extracellular signaling network and overview of the datasets considered. a) workflow of the procedure to generate enzyme-receptor interacting pairs; b) Sunburst charts providing an overview of the number of liganded and orphan G-protein-coupled receptors (GPCRs) considered; Classification of the liganded GPCRs based on their ligand type, i.e., either peptide, organic, or a combination of both; number of enzymes that are either currently included or absent within the Reactome pathways. The included ligands are additionally subcategorized based on the frequency of their types and the frequency of the Reactome pathway domains with which they are linked. For enzymes, only the second layer of this information is shown; c) The network of GPCRs and cognate ligands and biosynthetic enzymes; d) Funnel plot exhibiting the frequency of differentially expressed (TCGA vs. GTEx, BH corrected Padj<0.01) GPCRs, ligands, and enzymes corresponding to each whole tissue. e) Stacked bar-plot displaying the number of significantly enriched ligand-associated pathways within Reactome sub-domains: Signaling (‘R-HSA-372790: signaling by GPCR’) and Metabolism (‘R-HSA-1430728:metabolism’). Each bar represents the relative proportions of different categories as percentages of the whole, with respect to each tissue. An enriched pathway refers to a pathway enriched in TCGA with a GSEA enrichment score>0 and BH corrected Padj < 0.01.

We then computed genome-wide differential expression (DE) by contrasting cancer samples from the TCGA to those from corresponding healthy tissues from GTEx to derive expression signatures of genes and pathways mediating GPCR signaling. We considered a total of 15000 samples, of which 10500 were from TCGA and 4500 from GTEX, encompassing 16 tissues, as well as 79 cancer molecular subtypes (see Methods; Supplementary Table 3).

Remarkably, we found a widespread, cancer-specific regulation of GPCR-related genes with respect to healthy tissues. We found on average 91 up-regulated and 92 down-regulated GPCRs (FDR < 0.01, |log2FC|>1) across cancer tissues. When considered together with their cognate ligands and biosynthetic enzymes, we obtained an average of 161 up- regulated and 133 down-regulated instances (Figure 1d). The amount of significantly DE genes varies across tissues, with Testis, Skin and Ovary being the most affected, and Bladder the least (Figure 1d). Certain cancer tissues are characterized by a prevalence of either significantly over-expressed (e.g., Pancreas) or downregulated (e.g., Brain) GPCRs (Supplementary Figure 1a). We found no significant trend when grouping DE receptors based on their G-protein coupling specificity (Supplementary Figure 1a). On the other hand, DE profiling of heterotrimeric G-protein α subunits showed that *GNAS* is the single subunit most widely up-regulated, being significantly over-expressed in 94% of cancer tissues (Supplementary Figure 1b,c), confirming previous analyses^15,17^.

At the process level, we found a widespread DE regulation (FDR < 0.01) of GPCR pathways across cancers (Figure 1e). In particular, 40% of cancer tissues are either more enriched in biosynthetic or signal transduction processes, while 20% of cancer tissues are equally affected in both (Figure 1e). Several pathways are found to be widespread affected (e.g., up- regulation of Chemokine receptors), while others are exclusively regulated in certain tumor types (Supplementary Figure 1d). Overall, these data show that every cancer tissue is characterized by a highly specific GPCR pathway signatures (Supplementary Figure 1d).

### The landscape of peptide ligand-GPCRs dysregulation in cancer

We focused our analysis on the extracellular components of the GPCR signaling machinery, e.g., ligands and related metabolizing enzymes, to understand their contribution to the overall dysregulation of GPCR signaling in cancer.

We found on average 66 significantly DE peptide ligand precursors in 16 cancer tissues, and on average 60 instances in 79 molecular subtypes.

We next assessed the degree of correlation of expression change between receptors and interacting ligands and found a widespread co-regulation of receptor-ligand precursor pairs in multiple cancer tissues and subtypes (Figure 2a, Supplementary Figure 2; see Methods). By focusing on the receptor-ligand pairs more recurrently co-regulated, we identified two clusters, showing respectively a prevalence of concordantly up-regulated (i.e., Cluster 1) and concordantly down-regulated (i.e., Cluster 2) axes across subtypes (Figure 2a). The majority of the most frequently regulated ligands are agonists, with the exception of *CCL4*, *CCL7* and *CXCL8,10,11*, which might act as either agonists or antagonists depending on the bound receptor (Figure 2a). Most of axes in Cluster 1 involve chemokine receptors (e.g., *CCR1, CCR3, CCR4, CCR5, CCR6, CCR8, CXCR3, CXCR6, XCR1*) as well as other receptors such as *APLNR*, *CALCR*, *GRPR, MCHR1, GPR37, CCKBR, NPBWR1, KISS1R* and their cognate ligands. The main transduction mechanism in this group is through G_i/o_ proteins, as well as to a lesser extent via G_q/11_ and G_12/13_ (Figure 2a).

**Figure 2.**
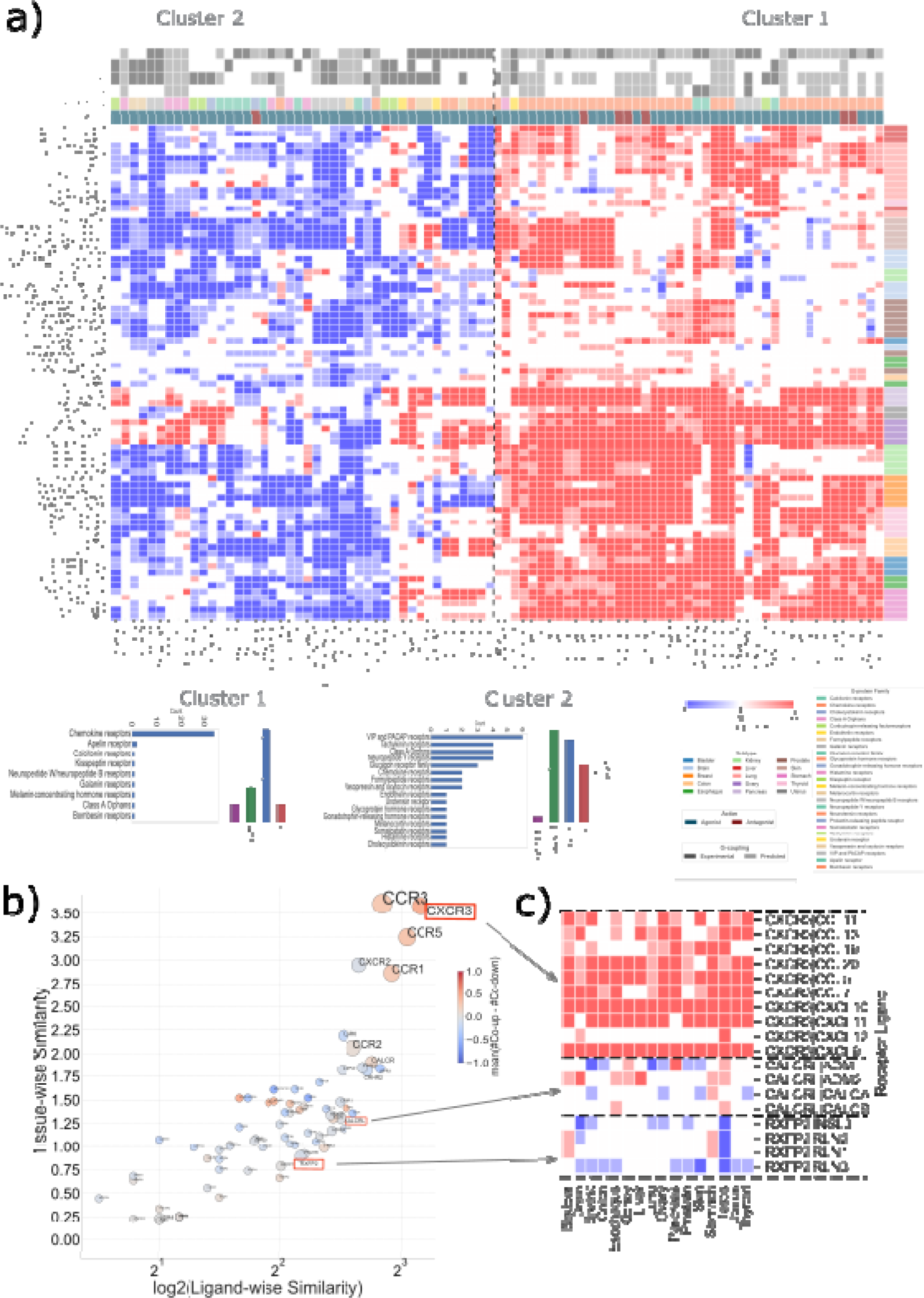
Peptide Ligand-Receptor co-regulation. a) Heatmap (center panel) displaying the co-differential regulation for Receptor-Ligand pairs across different TCGA subtypes (color- coded at the top row). Darker red represents both receptor and ligand significantly co-up regulated in TCGA (i.e., LFC>1 and Padj<0.01) and darker blue represents both receptor and ligand significantly co-down regulated in TCGA (i.e., LFC<1 and Padj<0.01). Paler colors represent either of the receptor-ligand as significantly DE (i.e., |LFC|>1 for both but Padj<0.01 for only one of these). White cells indicate anti-regulation or no fold change at all in at least one of them. Only those pairs which are affected in at least 25% of TCGA subtypes are displayed. A dashed line separating the two clusters, created using Hierarchical clustering, is shown in the middle. Hierarchical clustering was performed using the ’ward’ method to identify two homogeneous clusters by minimizing within-cluster variance based on the sum of squared differences of feature values i.e. scores∈[-2,+2] assigned to each pair. Heatmap (left panel) uses color codes to display the ligand’s mechanism of action in ‘Action’, G Protein-coupling associated with the GPCRs and GPCR Family. Heatmap (right panel) displays the information contained in the left panel as concise bar plots for each of the two clusters. For each cluster barplots indicate G-protein coupling preferences as well as GPCR families represented as count of the number of recepors; b) The scatter plot displays a GPCR-centric view of the heatmap in (a) based on tissue-wise or ligand-wise similarity. Here, ligand-wise similarity for each GPCR is the average of the pairwise Euclidean distances between co-differentially expressed (co-DE) profiles across its ligands (i.e., vertically), while tissue-wise similarity is a similar metric across tissues (i.e., horizontally). The size of the markers is proportional to the number of ligands, and the colors correspond to the average difference in the number of co-up and co-down profiles. GPCRs located in the top-right corner are more diverse in terms of their ligand interactions; c) Diverse co-DE profiles are observed with respect to different ligands and tissues corresponding to the same GPCR. For example, CXCR3 and its corresponding ligands exhibit an almost exclusively co-upregulated profile irrespective of tissue type, while CALCRL and its ligands display varied co-regulation profiles, implying a varying interaction with its ligands in different tissue (or cancer) types.

Cluster 2 contains instead receptor-ligand axes that are more frequently concordantly down- regulated in cancer subtypes, although several of these are concordantly up-regulated in other specific subtypes (e.g., *CXCR2, EDNRB, FFPR2*, or *LGR6, GALR2* and *UTS2R;* Figure 2a). The axes in Cluster 2 involve receptors that are members of various families from class A and B. Receptors in this group have a more heterogeneous/promiscuous coupling profile to G_q/11_, G_i/o_ and G_s_, but very little to G_12/13_ (Figure 2a).

In general, every cancer subtype is characterized by a highly specific receptor-ligand co-expression signature. However, certain tissues display similar subtype co-expression patterns (e.g., liver, breast, ovary, thyroid, stomach, lung, skin and pancreas). Others show higher heterogeneity (e.g., kidney, colon, bladder, esophagus, prostate, brain) (Figure 2a).

Co-regulation patterns of a given receptor can vary, depending on the tissue as well on the number and type of bound ligands (Figure 2b, Supplementary Figure 2). Certain receptors display consistent co-regulation patterns across cancer tissues and subtypes even if bound to multiple ligands (e.g., *CXCR3, CCR3*, *CCR5*; Figure 2b,c). Others show more variable co-regulation patterns which vary in a tissue (or subtype) and ligand specific fashion (e.g., *CALCRL* or *RXFP2*; Figure 2b,c).

### Expression of biosynthetic pathway enzymes inform on the levels of organic endogenous ligands

We reasoned that the expression levels of the enzymes synthesizing organic GPCR ligands could be informative of their levels and thus be used as a proxy to model ligand abundance in transcriptomic datasets. To this end, we assessed enzyme and biosynthetic pathway activities from differential expression signatures (see Methods).

We found on average 63 and 8 significantly regulated enzymes and biosynthetic pathways, respectively, across cancers, suggesting a widespread dysregulation of GPCR ligand biosynthetic pathways in human malignancies, which in some cases were specific to distinct cancer subtypes (Figure 3a). Clustering of biosynthetic pathways via enrichment scores revealed two main groups: one characterized by processes, such as those related to selenocysteine synthesis and di/triphosphate nucleotide interconversion, which are overall up-regulated in most cancer subtypes, while the others are characterized by processes recurrently down-regulated, such as “Phase I - Functionalization of compounds” or “Eicosanoids” (Figure 3a). Other biosynthetic pathways are specifically enriched in given cancer subtypes. For example, “Arachidonic acid metabolism” is down-regulated in several Kidney and Skin cancer subtypes, while it is significantly up-regulated in Ovary “Mesenchymal” and Glandular “GL” Pancreas subtypes. Other pancreatic cancer subtypes, such as the Transitional (“TR”) and Undifferentiated (“UN”), displayed instead significant enrichment of “Synthesis of Prostaglandins (PG) and Thromboxanes (TX)” pathway (Figure 3a).

**Figure 3:**
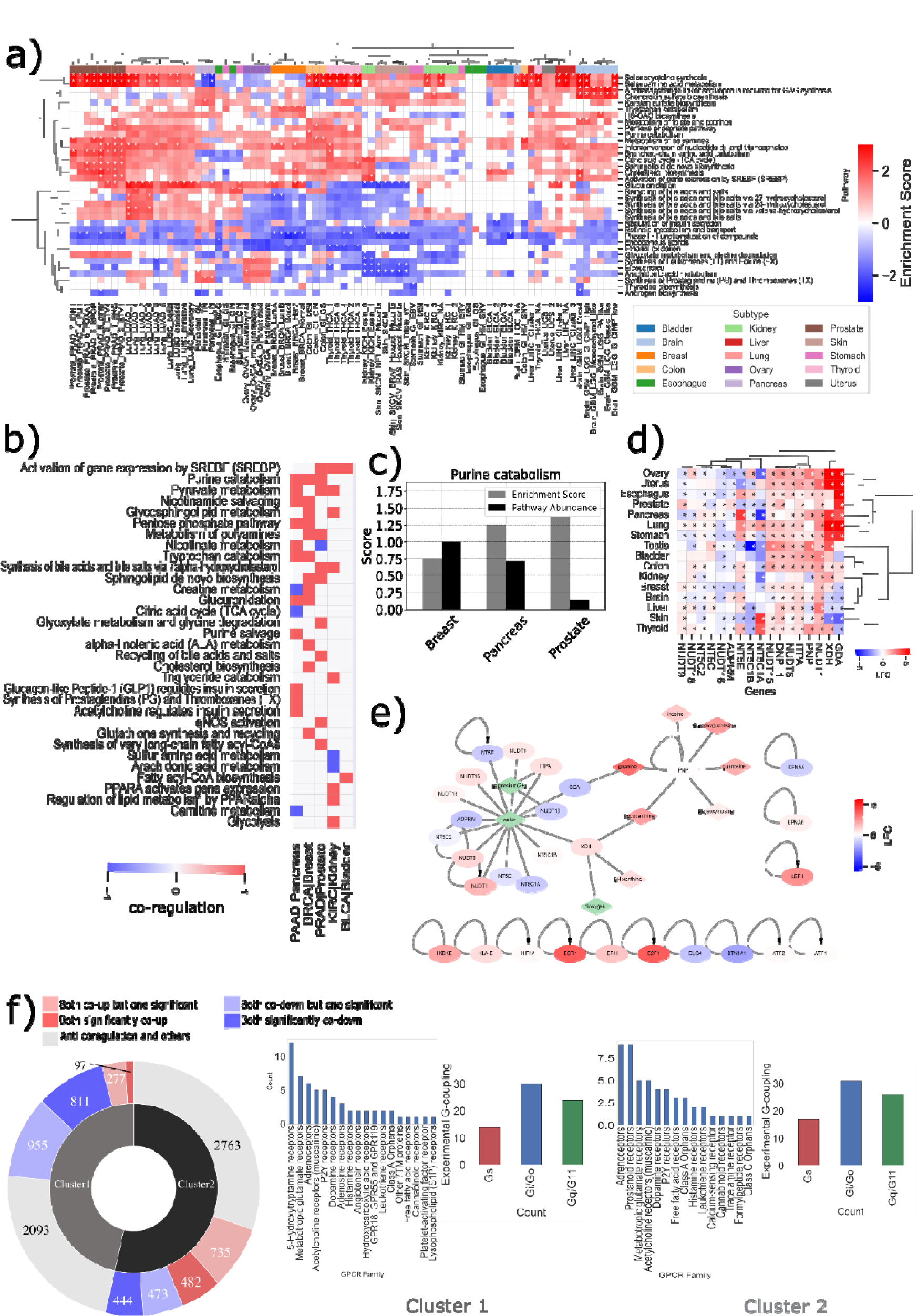
Biosynthetic pathway enrichment. (a) Heatmap of GSEA pathway enrichment analysis for differentially expressed (DE) genes in cancer (TCGA) over normal (GTEx) samples, considering Reactome pathways related to signaling and metabolism. Red cells indicate pathways enriched in TCGA, blue cells indicate pathways enriched in GTEx, and white cells indicate no enrichment. Cells marked with ‘*’ correspond to a statistically significant enrichment (Padj < 0.01). (b) Concordant pathway enrichment analysis considering both DE genes (Enrichment Score, ES) and metabolites (Pathway Abundance Score, PAS). A score of 1 (red) indicates co enrichment in cancer (ES & PAS > 0), and a score of -1 (blue) indicates co enrichment in normal samples (ES & PAS < 0). (c) Bar plot displaying numerical values for ES and PAS for the ‘Purine catabolism’ pathway in three different tissues: Breast, Bladder, and Prostate. (d) Heatmap of DE genes for the ‘Purine catabolism’ pathway in Breast cancer. Red cells indicate upregulated genes in TCGA, blue cells indicate downregulated genes in TCGA, and cells marked with ‘*’ correspond to a statistically significant differential expression (Padj < 0.01). (e) A functional interaction network between genes (ovals) and metabolites (diamonds) in the ‘Purine catabolism’ pathway in Breast cancer. Red nodes indicate upregulated components, blue nodes indicate downregulated components, and green nodes indicate no information available. The network shows that over-activation of the pathway is contributed by over-expression of both genes and metabolites; (f) Sunburst chart (left panel) displaying the distribution of co-differentially regulated Receptor-Enzyme pairs across two different clusters similar to receptor-ligand co-regulation analysis (Supplementary SX). Darker red represents the number of Receptor-Enzyme pairs significantly co-up regulated in TCGA (i.e., LFC>1 and Padj<0.01) and darker blue represents the number of Receptor-Enzyme pairs co-down regulated in TCGA (i.e., LFC<1 and Padj<0.01). Paler colors represent the number of pairs wherein either of the receptor-enzyme is significantly DE (i.e., |LFC|>1 for both but Padj<0.01 for only one of these). Grey indicates pairs with anti-regulation or no fold change at all in at least one of them. Only those pairs which are affected in at least 25% of TCGA subtypes are utilized for the clustering. For each cluster, barplots (center and right) indicate G-protein coupling preferences as well as GPCR families represented as count of the number of receptors

In order to determine if DE gene transcripts are mirrored by differential expression of the corresponding metabolites, we compared pathway enrichment scores of DE transcripts with differential abundance scores of DE metabolites from a pancancer metabolomic profiling dataset^24^, considering a set of curated biosynthetic pathways (e.g., Reactome; see Methods;Supplementary Table 4). We found a total of 33 significant pathways (FDR < 0.01) in at least one of the 5 cancer types considered, and which showed concordant pathway enrichment when considering both differentially expressed transcripts and metabolites (Figure 3b). The cancer type showing the highest number of concordant signatures is pancreatic adenocarcinoma (PAAD). Certain pathways show concordant up-regulation in multiple cancer types, including: “activation of gene expression by SREBF (SREBP)”, “Purine catabolism”, “Pyruvate metabolism”, each found up-regulated in three distinct cancer types (Figure 3b). For example, “Purine catabolism” is always up-regulated when considering both transcriptome and metabolome in Breast, Pancreas and Prostate cancers (Figure 3c). Inspection of differentially expressed genes and metabolites on a functional interaction network shows that over-activation of the pathway is contributed by over- expression of both transcripts and metabolites components (Figure 3e). However, only a few enzymes or metabolites in this pathway are invariably over-expressed across cancers (e.g., *NUDT1*, *LEF1*, 9H-Xanthine; Figure 3d, Supplementary Figure 3). Other processes show concordant enrichment with a cancer dependent directionality: e.g., nicotinate metabolism, which is up-regulated in pancreas and down-regulated in prostate, or creatine metabolism, down-regulated in pancreas and up-regulated in breast (Figure 3b). Several other pathways are exclusively enriched in specific cancer tissues: for example, “Glucagon-like Peptide-1 (GLP1) regulates insulin secretion”, “Synthesis of Prostaglandins (PG) and Thromboxanes (TX)” and “Acetylcholine regulates insulin secretion” are hallmarks of pancreatic cancer and might contribute to reinforce the known role of the Gs-PKA signaling axis in this cancer type ^25,26^.

Similarly to receptor-ligand pairs, we correlated changes in expression for receptor-enzyme pairs. Also in this case, we found two major clusters, respectively characterized by a majority of concordantly up- and down-regulated enzyme-receptor pairs (Figure 3F and Supplementary Figure 4). In particular, in the first cluster we found exclusive up-regulation of axes including those of 5-hydroxytriptamine and adenosine receptors with biosynthetic enzymes for their cognate ligands (Figure 3F and Supplementary Figure 4). In the second cluster, we found exclusive concordantly down-regulation of prostanoid and calcium-sensing receptors (Figure 3F and Supplementary Figure 4). Other receptor classes are represented in both clusters, via the involvement of distinct receptors and enzymes members (e.g. Adrenoceptors, Metabotropic glutamate receptors). On average, we found 63 receptor- enzyme pairs whose expression is significantly correlated with respect to a background random model (Supplementary Figure 5, Supplementary Table 5; see Methods).

### Expression of GPCRs signaling network components correlates with patient survival

We systematically assessed correlations between expression of GPCR network components and patient survival. Briefly, we defined groups of patients based on the expression of a GPCR component, i.e., high or low expression, respectively, if above or below the median of the distribution of expression values of that gene in the samples and tested whether these groups had significantly different survivals (see Methods).

We first considered the capability of the expression of individual components (e.g., receptors, transducers, ligand or biosynthetic enzymes) to discriminate patients based on their survival. We found a total of 302 unique significant receptor instances (LogRank P-value < 0.05, FDR < 0.1) in 11 cancer tissues, 103 ligands in 10 cancers, and 71 enzymes in 10 cancers (Figure 4a; Supplementary Table 6-8). Some cancer tissues display overall more components whose expression is associated with higher (e.g., pancreas, skin or head and neck) or lower (e.g., brain, breast, lining of body cavities and white blood cell) survival (Figure 4a). Overall, the expression of GPCR genes is associated with poorer survival in the 59% of cases across all cancers. *CELSR3*, *GPR25, EDNRB and F2RL2* are significantly associated with patients’ survival in four cancers each, with an equal prevalence for lower and higher survival (Figure 4b, Supplementary Table 6). Some receptors show consistent associations across cancers: for example, *ADORA2A* is invariably associated to higher survival across four distinct cancer tissues (including pancreas, breast, skin and head and neck; Figure 4b), which might be consistent with its known role in regulating CD8+ T cell activity and survival in the tumor microenvironment^27^. Certain receptors are instead invariably associated with poorer survival, such as *OXTR, ADORA2B, GPR3, FZD6, GPRC5A and LPAR3*. We also found a large prevalence of associations between GPCR peptide ligand precursor expression and lower survival of patients, with 67.2% of the significant instances across cancers. Among the most recurring ligands, we found several instances that are either invariably associated to lower survival, e.g., *CXCL5*, *CXCL1, GAL* and *RPS19*, or have opposite associations depending on the cancer type, e.g., *CXCL9, C3, CCL26, CCL8, CCL13, RSPO1, RLN1, RLN2* and *UCN* (Figure 4b; Supplementary Table 7).

**Figure 4.**
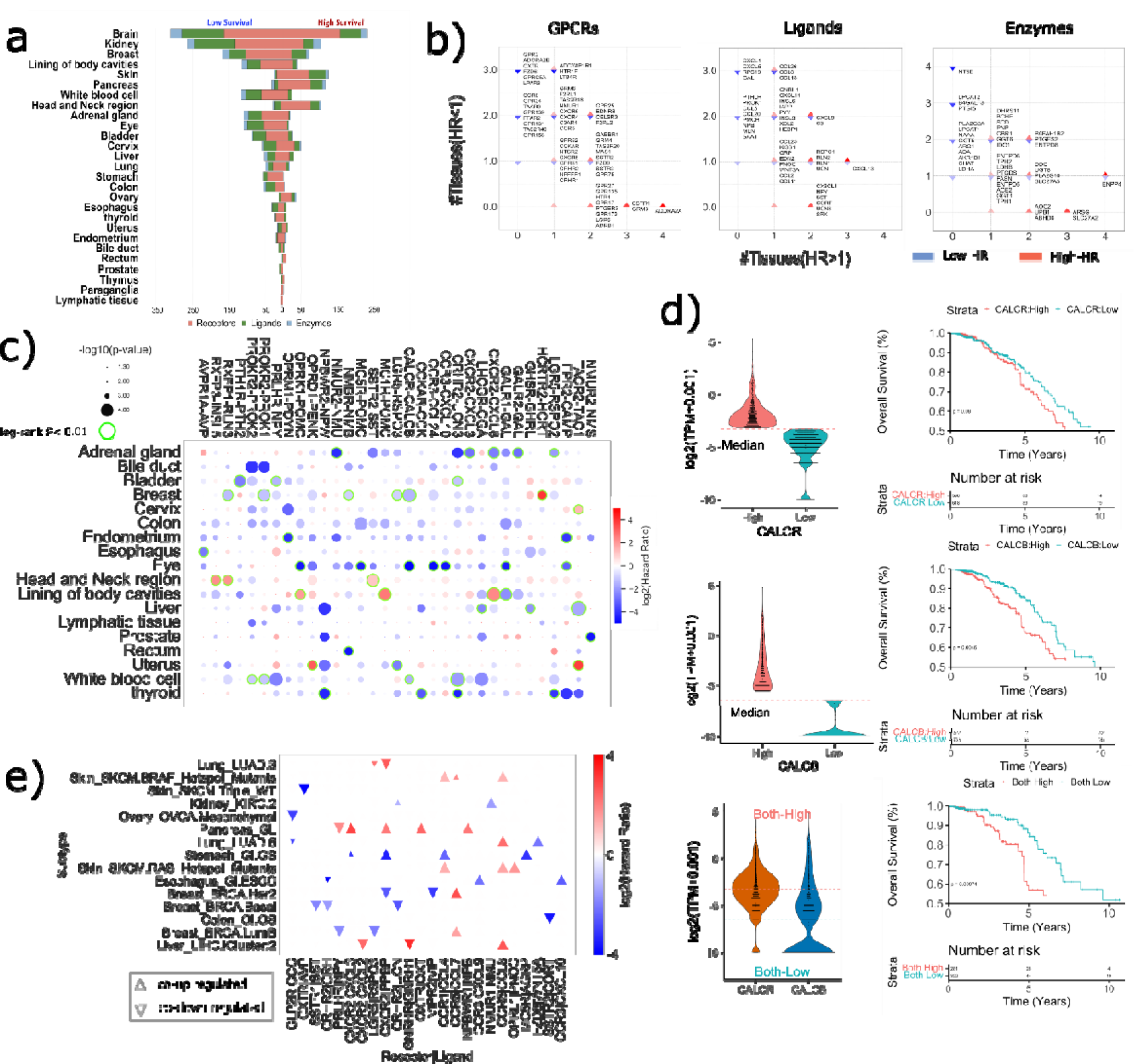
Association of GPCR-peptide ligands axes to survival. (a) Survival association of G protein-coupled receptor (GPCR) components in various cancer types. The funnel plot shows the number of significant instances of individual components, such as receptors, ligands, and enzymes, whose expression values are associated with patient survival across various cancer types. A total of 302 unique significant receptor instances (log rank p-value < 0.05, FDR<0.1) were identified in 11 cancer tissues, 103 ligands in 10 cancers, and 71 enzymes in 10 cancers. (b) Scatterplots for individual receptor instances, enzymes and ligands associated with patient survival in various cancer types. (logrank-p<0.05, FDR<0.1). x-axis displays the number of tissues in which the genes are associated with lower survival (inverted blue triangles) and, vice-versa, y-axis displays the number of tissues in which the genes are associated with higher survival (red triangles). Only the genes which are significantly associated with survival in at-least two tissues are shown. c) The bubble plot shows the correlation between the combined-expression levels of GPCR-ligand pairs and patient survival across various cancer types. The pairs with a log rank p-value <0.05 and also lower than log rank p-values for individual GPCR/ligand are displayed. Bubble color is proportional to HR: i.e., HR>1 : High expression is correlated with high survival (red); HR<1: High expression is correlated with poor survival (blue). Bubble diameters are proportional to the -log10(log-rank p-value). Green highlighted bubbles represent the most significant instances (sample sizes>5, FDR<0.1). (d) The expression values for the CALCR-CALCB axes in breast cancer (right) and corresponding Kaplan-Meier curve for survival analysis of patients stratified based on the individual as well as combined receptor/ligand expression;(e) Scatter-plot showing with differentially co-expressed GPCR-ligand pairs (upper triangle=co- up regulation; lower triangle=co-down regulation) that were significantly associated with patient survival (red=higher survival; blue=lower survival). A total of 24 receptor-ligand pairs from 15 cancer subtypes were identified.

We reasoned that in those patients with higher expression of both receptors and ligands (or biosynthetic enzymes), a likely enhanced activation of the signaling axis should better correlate with specific patient phenotypes (e.g., lower or higher survival) than when considering individual components. We therefore assessed differences in the survival of patients grouped on the basis of the expression levels of GPCRs and ligand precursors (or enzymes), either alone or in combination. We then shortlisted receptor ligand pairs characterized by higher significance as well as greater hazard ratios of patient survival compared to the individual interaction partners (Figure 4c; Supplementary Table 9-10, see Methods). Intriguingly, we found several axes, such as *NPBWR2-NPW, PRHLR-NPY, PRLHR-PLR, CRHR2-UCN3, CALCR-CALCB,* associated with lower survival more significantly than considering interaction partners alone in multiple cancer types (Figure 4c,d; Supplementary Figure 6). Certain cancer types show multiple significant instances of associations between receptor-ligand axis and survival (e.g., lower survival in adrenal gland, eye; Figure 4c), suggesting a potential polypharmacology strategy for these tumor types.

Interestingly, it appears that some receptors participating in significant axes within the same cancer type share, to some extent, similar signal transduction mechanisms, suggesting that the patient survival phenotype might be contributed by multiple ligand stimuli converging on common signaling cascades. For example, in head and neck cancers we found significant correlation of higher survival and higher expression of *SSTR2-SST* (Supplementary Figure 7a*), RXFP3-INSL5* and *RXFP1-RLN3* signaling axis, which all share similar coupling profiles (e.g., G_q/11_, particularly *GNA14*, as well as G_i/o_; Supplementary Figure 8). *RXFP1-RLN3* is instead associated with lower survival in breast carcinoma patients, highlighting the specificity of survival associations of these axes with cancer subtypes. Likewise, *SSTR2- SST* is associated to lower survival in Adrenal Gland patients (Supplementary Figure 7b), *OPRK1-POMC* with lower survival in uveal melanoma and higher survival in Lining of body cavities, *CXCR2-CXCL6* with lower survival in Adrenal Gland and higher survival in Lining of body cavities, *TACR2-TAC1* is associated with lower survival in Liver cancer and higher survival in Cervix and Uterus.

Notably, some of the receptor-ligand pairs that we found differentially co-regulated in terms of expression, are also significantly associated with survival differences (Figure 4e). In details, we found a total of 24 receptor-ligand pairs from 15 cancer subtypes. *CCR5* is the receptor-ligand forming pair (with *CCL7* and *CCL8*) that is more frequently concordantly DE- regulated, being at the same time significantly associated with survival. In particular, *CCR5* concordant up-regulation is invariably associated with a better prognosis, irrespective of the cancer subtype considered. Other pairs, such as those mediated by *CXCR2,* display cancer subtype-specific patterns of both differential expression regulation and correlation with patient survival (Figure 4e). On the other hand, pairs such as *CCR1-CCL4* are always co-up regulated but with different prognostic values, i.e., they are associated to higher survival in melanoma (RAS and BRAF hotspot mutants subtypes), and to lower survival in gastric cancer (“GI.GS” subtype, Figure 4e). The latter is also an example of cancer subtype with multiple axes association to lower survival. Other subtypes, such as the Glandular (“GL”) Pancreas, are instead characterized by multiple axes associated to higher survival (Figure 4e). Overall, we found 124 receptor-ligand pairs in 48 cancer subtypes more significantly correlated to survival than their individual instances (Supplementary Figure 9).

*Biosynthetic enzymes and cognate GPCRs form surrogate signaling pairs with prognostic value*.

We found multiple ligand-synthesizing enzymes whose expression correlates with patients’ survival in multiple cancers, with 70% of them significantly linked to worse survival (LogRank P-value < 0.05, FDR<0.1). Remarkably, the 5’-nucleotidase *NT5E* is invariably associated to lower survival in four distinct cancer types, including breast, brain, stomach and pancreas (Figure 4b; Supplementary Table 8). Other enzymes most recurrently associated with lower survival are Lysophosphatidylcholine acyltransferase 2 (*LPCAT2*), Beta-1,4- galactosyltransferase 3 (*B4GALT3),* as well as prostacyclin synthase (*PTGIS*). On the other hand, enzymes such as Bis(5’-adenosyl)-triphosphatase (*ENPP4)*, Arylsulfatase G (*ARSG)* and the Long-chain fatty acid transport protein 2 *(SLC27A2)* are recurrently associated to higher survival (Figure 4b).

We similarly checked for an improvement of the survival statistics when considering the combined expressions of receptor-enzyme interaction partners (Methods; Supplementary Table 10). This revealed several instances involving biosynthetic enzymes for neurotransmitters and cognate receptors (Figure 5A, Supplementary Table 10). Multiple pairs involve enzymes for monoamine neurotransmitter biosynthesis, such as Tryptophan 5- hydroxylase 2 (*TPH2*), the rate-limiting enzyme for serotonin synthesis, which is significantly associated to lower survival with multiple serotonergic receptors (e.g., *HTR1A* in Liver, *HTR2C* and *HTR6* in Adrenal Gland and *HTR7* in White Blood cell). Several Dopamine receptors are significantly correlated along with enzymes mediating catecholamine biosynthesis, including *DDR3-DDC* (liver), *DDR2* and *DDR3-TH* (prostate). The latter enzyme, which rate-limits the catecholamine pathway, is also significantly associated with *ADRA1D* in Lymphatic Tissue. The axes formed by Choline O-acetyltransferase (*CHAT)* and muscarinic receptors *CHRM1,3,5* are associated with lower survival in multiple cancers (Figure 5a, b). The three axes involving muscarinic receptors and *CHAT* are all associated to lower survival in liver cancer, with *CHRM3-CHAT* being also significantly correlated in Esophageal cancer (Figure 5a). We found multiple instances of glutamate receptors (e.g., *GRM1,2,4,5*) and Glutamate decarboxylase 1 and 2 (*GAD1,2*). While the *GRM2-GAD1* receptor-biosynthetic pathway pair shows tissue specific correlation with survival, being associated with lower survival in Adrenal Gland and with higher survival in Liver, the axes of *GAD2* with *GRM1,4,5* are always significantly associated with lower survival in Endometrium and Thyroid cancer tissues. We also found several instances of receptors for purines whose aberrant expression co-occur with that of multiple enzymes of the Purine catabolism pathway. In details, while the *ADORA2B-PNP* pair is associated to lower survival in Uterus (Figure 5a,b), *P2RY4* axes with *ENPP3* and *ENTPD1* are associated to higher survival in the same cancer. The latter axis is instead significantly associated with lower survival in Adrenal Gland (Figure 5a). Overall, we found 115 receptor-enzyme pairs in 45 cancer subtypes more significantly correlated to survival than individual instances (Supplementary Figure 10).

**Figure 5.**
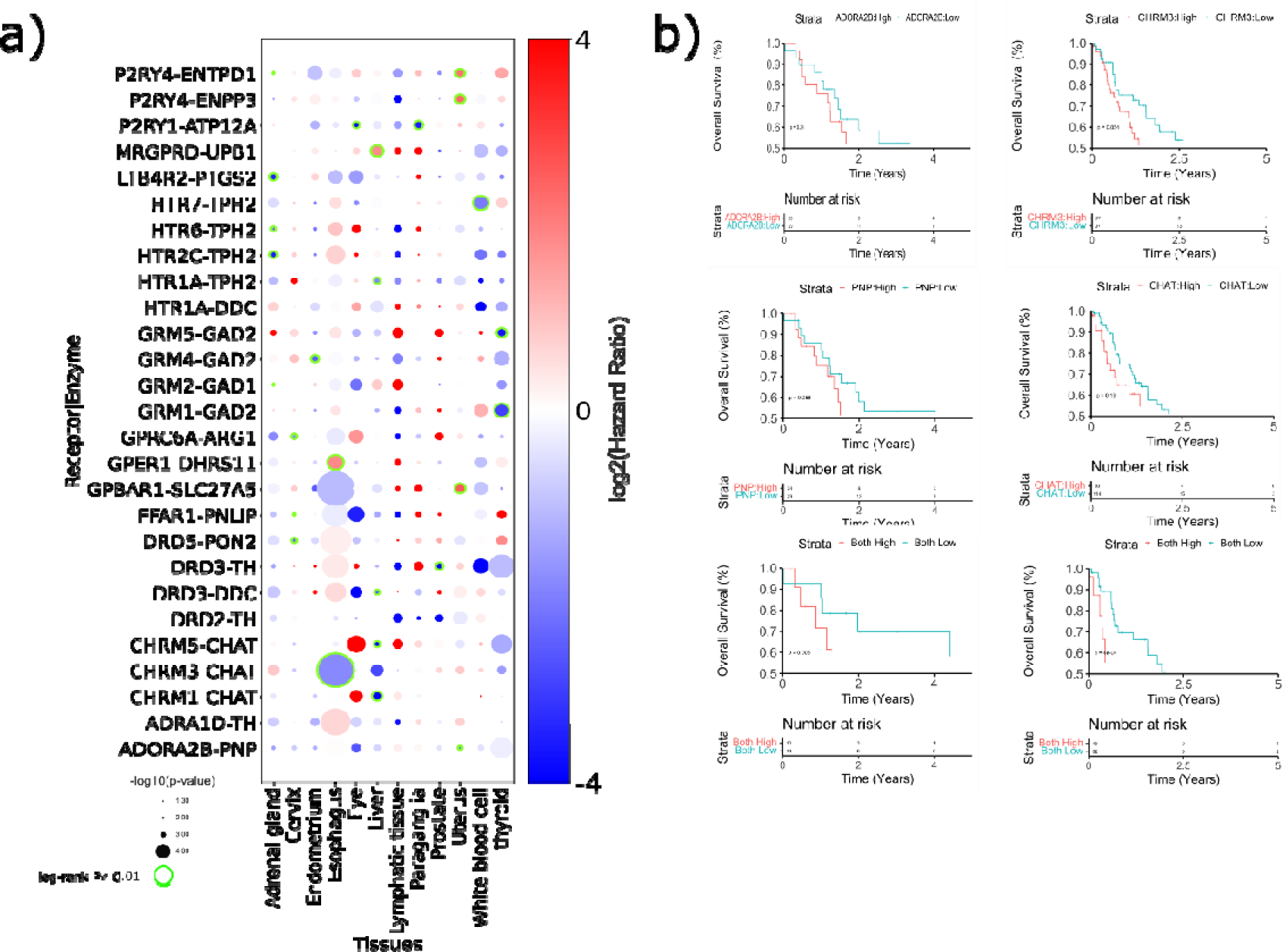
Association of GPCR-enzymes axes to survival. (a) The bubble plot shows the correlation between the combined-expression levels of GPCR-enzyme pairs and patient survival across various cancer types. The pairs with a log rank p-value <0.05 and also lower than log rank p-values for individual GPCR/enzyme are displayed. Further, the pairs with rate-limiting enzymes having less than 100 PubMed references are not shown. Bubble color is proportional to HR: i.e., HR>1 : High expression is correlated with high survival (red); HR<1: High expression is correlated with poor survival (blue). Bubble diameters are proportional to the -log(log-rank p-value). Green highlighted bubbles represent the most significant instances (sample sizes>5, FDR<0.1). (b) The CHRM3-CHAT axis is recurrently correlated with lower survival rates in Esophageal cancer. The KM plots (left) display the risk enhancement achieved when CHRM3-CHAT combined stratification was applied as compared to individual CHRM3/CHAT as evident by log rank p-values. Similarly, the ADORA2B-PNP axis is also consistently associated with lower survival rates in Uterine cancer, as evident from the KM plot on the right.

### Drugs for GPCR networks hold the potential to inhibits cancer cell growth

We retrieved GPCRs ligands showing cancer growth inhibitory capacity from a public resource containing the growth-inhibitory activity of 4,518 drugs tested across 578 human cancer cell lines^28^. A total of 13 out of 52 GPCR ligands tested were found to significantly inhibit the growth of cancer cell lines (Figure 6a)^28^. We checked the targets of these ligands on the pooled list of GPCRs that we found to mediate axes significantly associated with cancer survival. Strikingly, all these drugs bind to receptors that are significantly coregulated axes with either their ligands or biosynthetic enzymes (Figure 6a,b; Supplementary Table 11). In particular, among them we found multiple ligands targeting certain GPCRs classes, including adenosine receptors (i.e., CGS-15943, MRS-1220, SCH-58261), muscarinic receptors (i.e., VU0238429, xanomeline, terfenadine) and opioid receptors (i.e., BNTX and JTC-801). Interestingly, the vast majority (76%) of these drugs are antagonists. Moreover, receptors targeted by antagonists appear to be more promiscuous, and coupling primarily to G_q/11_ or G_i/o_, but also to a lesser extent to G_s_ and G_12/13_ proteins (Figure 6c).

**Figure 6.**
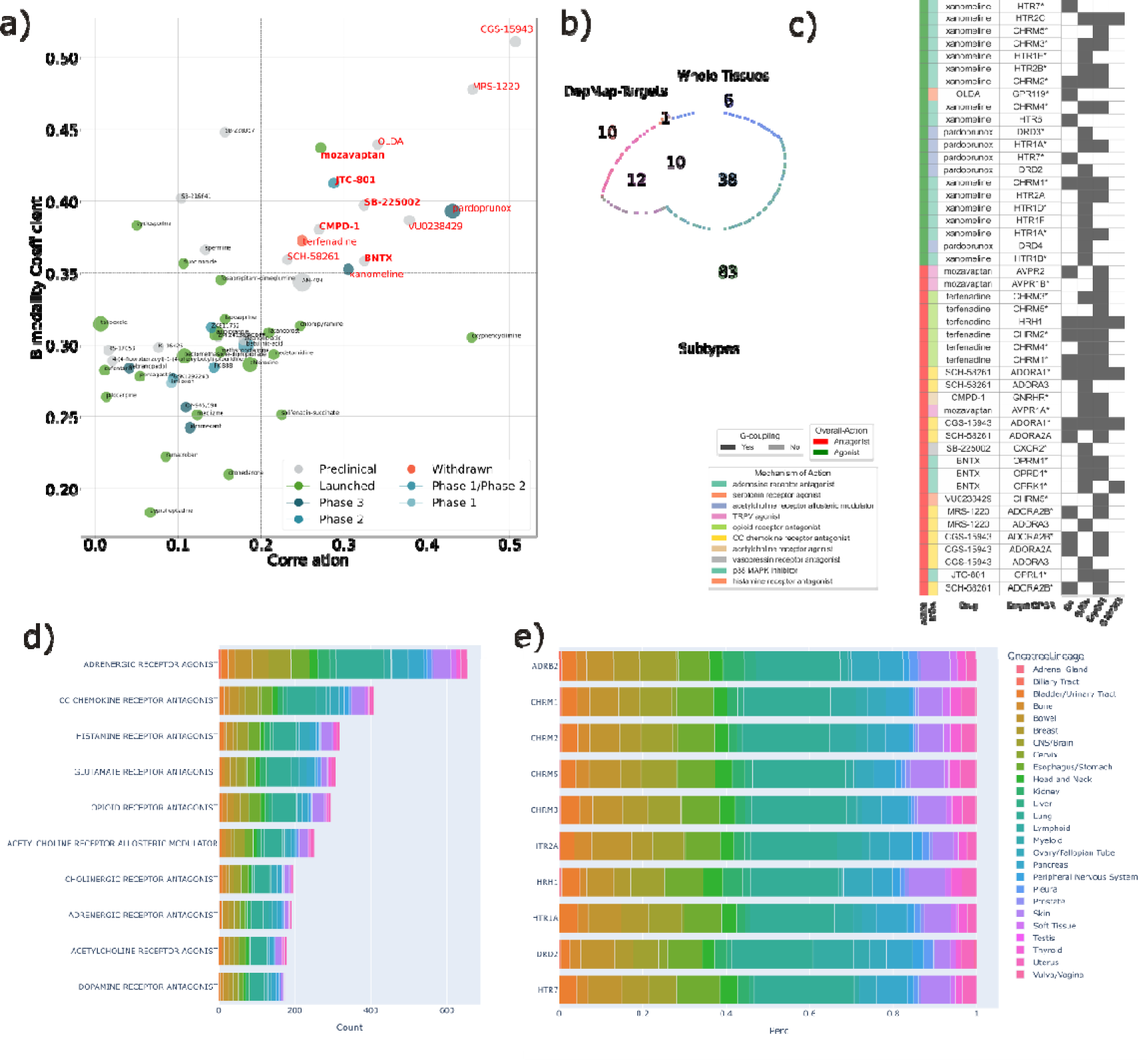
GPCR ligands with cancer cell line growth-inhibitory capacity. (a) The scatterplot displays 52 GPCR ligands that were tested in 578 cell lines in PRISM drug repurposing resource.13 GPCR ligands (in red) were found to significantly inhibit the growth of cancer cell lines (correlation>0.2, bimodality coefficient>0.35). (b) Venn diagram showing the overlap of GPCRs in the target proteins of these 13 ligands with the pooled list of GPCRs that were found to mediate survival associated GPCR-ligand/GPCR-enzyme axes. (c) The heatmap displays the 13 shortlisted ligands and their corresponding GPCR targets. The targets marked with ‘*’ are the overlapping GPCRs with significant axes. Heatmap also displays the overall action mode of ligand, more detailed mechanism of action (MOA) and G- protein coupling associated with the GPCR; d) stacked barplot of top 10 mechanisms of actions (MOA) involving the top 10 GPCR drug with the highest growth inhibition potential (ranked according to the negative of the log2fold change in each cell line). Each stack is proportional to the number of cell lines of a given tissue where a drug belonging to that MOA is ranked among the top 10 inhibiting drugs. Stack coloring is tissue specific; e) stacked barplot, normalized for the total cell line-drug pairs count, of top 10 drug targets hit by the top 10 GPCR drug with the highest growth inhibition potential (ranked according to the negative of the log2fold change in each cell line). Each stack is proportional to the number of cell lines of a given tissue where a drug hitting the specific target is ranked among the top 10 inhibiting drugs. Stack coloring is tissue specific.

We also considered an updated version of the PRISM assay of cancer cell sensitivity by pooling the data of 1514 GPCR drugs tested against 906 cell lines (Figure 6d,e; see Methods). The GPCR drugs with the highest potential of cancer cell line growth inhibition target adrenergic receptors, CC Chemokine, Glutamate, Histamine, Opioid, Acetylcholine and Dopamine receptors (Figure 6d). Among top ten individual targets is *ADRB2,* muscarinic, 5-hydroxytryptamine and *DRD2* receptors (Figure 6e). While cell lines from certain tissue seem to be equally inhibited by top GPCR drugs (e.g. lung), others are sensitive to specific GPCR drugs, such as Lymphoid cancer cells, which are particularly inhibited by *DRD2, CHRM1, CHRM2, HTR1A* and *HTR2A* antagonists (Figure 6e, Supplementary Table 12).

## Discussion

In recent years the multifaceted role of GPCR signaling in cancer has begun to emerge^15,16^, though an overall mechanistic picture is still lagging. At a molecular level, this is mainly due to the lack of a precise mechanistic understanding of how receptors transduce external signals intracellularly in the context of the TME, as well as how the upstream ligand and biosynthetic pathway components contribute to trigger an aberrant GPCR signaling network in tumor cells and their associated stroma and infiltrating immune cells. At a higher level, the complex circuitry wired by multi-component GPCRs systems is highly tailored to specific cellular contexts. Pathways dysregulation might be initiated through perturbations of these multi-component systems at different levels, all of which may converge to an overall dysregulation and/or persistent activation of GPCR downstream signaling.

In the present study, we systematically inferred GPCR pathways activities from cancer transcriptomic datasets, with a special focus on the dysregulation of GPCR signaling axes involving the co-deregulation of the expression of their direct ligands as well as biosynthetic pathways controlling their ligand availability.

We found that multiple GPCRs are co-regulated together with their cognate ligands or biosynthetic enzymes, suggesting a coordinated effect on the signaling axis. Some of these are widespread co-regulated across cancers, others are affected in specific tumor contexts. These studies revealed some exciting emerging patterns, among them that chemokine receptors and their cognate chemokines are certainly the most co-regulated axes. The precise role in pro- or anti-tumor immunity played by these chemokine receptor networks is known to depend on the precise cellular and tumor subtype context^15,29^. We have reported *CXCR3* to be concordantly up-regulated together with its ligands across cancer subtypes. Intriguingly, its expression in Treg cells has been recently shown to central for immune suppression of the CD8+ -mediated response in multiple cancers^30^. The known tumor-promoting *CXCR2* is found co-down regulated together with *CXCL2* in liver cancer (“iCluster2” subtype) and with *PPBP*, the precursor of *CXCL7*, in lung (“LUAD.3” subtype). Remarkably, in both cases, the co-down-regulation is associated with higher survival (Figure 4e). The co-up regulation of this axis is associated with lower survival in stomach cancer (“GI.GS” subtype) and with higher survival in pancreatic cancer (“GL” subtype), similarly to *CXCR2*-*CXCL3*. Other chemokine receptors, such as *CCR1, CCR3* or *CCR5*, are significantly co-up regulated across different cancer subtypes. While *CCR1-CCL4* has distinct correlation to survival depending on the cancer type considered, *CCR5-CCL7* or *CCR5-CCL8* co-up regulation is always associated with higher survival, suggesting a general anti-tumor immunity of these axes in the cancer subtypes considered (Figure 4e).

Interestingly, we also found a few non-chemokine receptors that are widespread co-up activated, such as *KISS1, NPBWR, MCHR1 or APLNR* which suggest the existence of signaling mechanisms other than chemokine sustaining the cell-cell interaction within the TME. Another notable example of highly context specific co-regulation of signaling axes is the one mediated by *CALCR* and its close paralogue, *CALCRL*, whose interaction with *ADM* is a critical factor determining relapse and drug resistance in Leukemia^31^. We found that many of the co-regulated signaling axes are also associated with patient survival. For example, in head and neck cancer we have found multiple signaling axes significantly associated to higher survival, such as *SSTR2-SST*, *RXFP3-INSL5* and *RXFP1-RLN3*. Intriguingly, all the three receptors show converging couplings towards G_q/11_ and Gi/o proteins. Notably, the *SSTR2-SST* has been significantly associated to higher survival rates in EBV-nasopharyngeal cancer, a class of head and neck cancer, and its targeting with a miniaturized peptide-drug conjugate (i.e., PEN-221) has been proposed to treat this type of tumor^32^. *SSTR2-SST* is also one of the axes with the strongest literature support to a role in cancer, along with several others including e.g., *MC1R-POMC*, *GHRHR-GHRH* (Supplementary Figure 11; Supplementary Table 13).

Along with the mRNA expression of peptide ligand precursors, we also considered the expression of synthesizing enzymes as a proxy to model upstream signals from small organic ligands. Indeed we found a significant correlation of patients’ survival with expression of axes formed by receptors and biosynthetic enzyme pairs. Similarly to the receptor-ligand axes, we found receptor-enzymatic axes recurrently associated with survival in a cancer-type specific fashion. For example, we found multiple instances of muscarinic receptors and Choline O-acetyltransferase (*CHAT*) co-regulation with patient outcome. Cholinergic signaling has been established as a key factor mediating gastric cancer tumorigenesis via an NGF promoted feed-forward loop^20^. Intriguingly, the association between the expression of muscarinic receptors-*CHAT* axis with lower survival in hepatocellular carcinoma correlates with the known involvement of acetycholinesterase, which opposes *CHAT* function by degrading acetylcholine, and is associated to better prognosis in HCC^33^. Other biosynthetic enzyme axes that we have identified are also supported by additional lines of evidence from the literature (Supplementary Figure 11d), and strongly emphasize the role of several neurotransmitter signaling in fostering oncogenic growth. The information about biosynthetic enzymes is indeed particularly important for receptor systems such as GPCRs, which frequently bind to organic ligands, and might represent a resource complementing existing knowledge-bases for receptor-ligand interactions used to characterize cell-cell communication from spatial transcriptomics datasets^23^. It will be critical in the future to dissect the precise cell context-specific role played by these GPCRs signaling axes by analyzing single cell as well as spatial transcriptomics RNAseq datasets, as we recently began exploring in the context of CD8-T cell function^34^.

Taken together, we have found a few hundreds of receptor pairs with either ligands or enzymes to be significantly associated to survival in tens of cancer molecular subtypes, suggesting a host of actionable opportunities to modulate oncogenic phenotypes. Strikingly, all the GPCR drugs that have been reported to significantly hamper cancer cell line growth in a recent systematic screen^28^, indeed target receptors of axes that we found significantly correlated with patient survival, with a particular emphasis on receptors for neurotransmitters and adenosine. In addition to directly hampering cancer growth, the targeting of GPCRs network is emerging as a promising therapeutic option in combination with immunotherapy^15^. For instance, adenosine signaling is an established factor which dampens the immune response in inflamed tissues and targeting of either adenosine receptors (particularly *ADORA2A*), or adenosine synthetizing enzymes (*NT5E* and *ENTPD1*) are currently being evaluated in combination with cancer immunotherapies^35,36^. Moreover, the G_s_-PKA axis has been recently identified as a key pathway dampening the anti-tumor CD8 T cell activity and resulting in Immune Checkpoint Blockade failure. This study suggests that additional G_s_PCRs (e.g., EP_2_, EP_4_, β_1_AR, and β_2_AR), other than *ADORA2A*, could be targeted in combination with immunotherapy^34^.

Ultimately, our data-driven approach, that integrates high-dimensional transcriptomics datasets and highly curated signaling networks, may enable the functional interpretation of GPCRs’ dysregulation mechanisms and guide the development of novel pharmacological strategies and drug repurposing opportunities targeting GPCRs in cancer.

## Methods

### GPCR signaling network

To create an extended GPCR network, we have extracted the GPCR and ligand interactions from the IUPHAR/BPS Guide to PHARMACOLOGY database (IUPHAR database)^37^. We retained only endogenous ligands, thereby extracting 243 GPCRs and 328 ligands. Based on database classification, the considered ligands can be grouped into five groups: 211 peptides, 107 metabolites, 7 synthetic organic, 2 inorganic and 1 natural product. All the other annotations describing GPCRs and ligands were also pulled from the database. To further extend our network we added biosynthetic enzymes for GPCR endogenous ligands. To this end we used Rhea, a curated database of chemical reactions of biological relevance^38^. To identify metabolizing enzymes for GPCRs ligands, we first referenced IUPHAR ligands to ChEBI^39^, which is used by Rhea to identify ligands. Ligand mapping between IUPHAR and Rhea was carried out by using the IUPHAR International Chemical Identifiers (InChIs). As many of the chemical reactions are happening under physiological pH of 7.3, the ChEBI identities of participants in Rhea reactions at pH 7.3 were considered. Using our in-house python scripts, we have extracted all the enzymes catalyzing reactions where GPCR ligands participate as products, meaning that most of the reactions have left-right (LR) directionality or, in some cases, opposite (right-left - RL) direction. Moreover, we also considered all the enzymes catalyzing reactions of unknown (UN) direction. Since it is possible that enzyme annotation is available only for parent reactions, i.e., more generic chemical reactions, we also added parent reactions based on Rhea’s hierarchical reaction relationships. For each of the Rhea identities acquired, we extracted the corresponding human enzyme whenever available. We extracted 1492 enzymes linked to GPCR ligands, which were filtered according to the following Gene Ontology cellular components terms: *’plasma membrane’, ’axon terminus’, ’external side of plasma membrane’, ’extracellular’, ’neuron projection’, ’channel’, ’membrane transport’*. We also included in our composite filter additional annotation terms from Pfam^40^, Panther classification^41^, Gene ontology processes (GO)^42^ or full UniProt^43^ enzyme name to filter out enzymes classified as kinases, channels, transporters, cytochromes and adenylyl cyclases. We also included rate-limiting enzymes, by mapping 3k human enzymes from RheaDB onto Reactome pathways from ’Metabolism’ domain (R-HSA-1430728), which were first filtered, using similar class and domain annotations as above, to exclude kinases, channels, transporters, cytochromes and adenylyl cyclases resulting in a total of 2261 unique enzymes, out of which 945 were found to match 163 pathways involving GPCR ligands. To determine which of these enzymes were rate-limiting the whole biosynthetic pathway, we retrieved for each of them the number of PubMed articles using the following search query: ’{Enzyme gene symbol} AND (rate limiting OR bottleneck)’. The enzyme with the highest number of PubMed references for each metabolism pathway associated to GPCR ligands was retained (Supplementary Table 14).

After these filtering steps, we yielded a total list of 266 enzymes, each involved in interaction with one or multiple GPCRs mediated by synthesized ligands. Receptor-enzymes were further filtered out by using the STRING database, a knowledgebase of known or predicted protein-protein functional interactions^44^. Via STRING, we mapped all possible interactions between GPCRs and enzymes producing cognate ligands, by using a minimum confidence score of 150, and leading to a shortlist of 82 enzymes. The final network contained 82 enzymes, 328 ligands and 243 receptors, for a total of 150 binary enzymes-ligand interactions, 646 ligand-GPCR, and 288 receptor-enzyme (ligand-mediated) interactions.

The whole network has been compiled containing different descriptions of each of the three main interactors and their interactions are currently being deposited to SIGNOR^45^ database thanks to an ongoing dedicated curation project.

We generated G protein specific gene set as well as annotations by gathering consensus experimental transducer data (i.e., Universal Coupling Map^9^) as well as predicted through Precogx^7^.

### Transcriptomics Dataset

We considered gene-level expressions from the dataset ‘TCGA TARGET GTEx’ publicly available at UCSC Xena (https://xenabrowser.net/datapages/)^46^. These data were generated using the TOIL pipeline, a portable, open-source workflow software that can be used to run scientific workflows on a large scale in cloud or high-performance computing (HPC) environments^47^. The Toil pipeline uses STAR^48^ to generate alignments and read coverage graphs, and performs quantification using RSEM^49^ and Kallisto^50^. It was utilized to process ∼20,000 RNA-seq samples to create a consistent meta-analysis of datasets free of computational batch effects.

We extracted the gene-level expression data, *RSEM expected_count*, using the R package ‘UCSCXenatools’ for 19109 samples. Thereafter, the TARGET samples were filtered and only TCGA and GTEx samples were retained (n=18305; Supplementary Table 2). Due to unavailability of GTEx data for all the tissue types, only 16 whole cancer-types (TCGA) were contrasted to the healthy samples pertaining to the same tissue (GTEx). The information for 76 distinct TCGA subtypes was retrieved from TCGAbiolinks^51^ (R4.2, Bioconductor v3.16) except for ‘Pancreas’ and ‘Testis’ for whom it was unavailable.

For pancreatic cancer we employed a new classification of TCGA samples using a multi-class classifier already implemented to classify three novel PDAC subtypes (Glandular, GL; Transitional, TR; Undifferentiated, UN) obtained from the transcriptional profiles of multiple morphological distinguishable tumor areas isolated by laser micro-dissection (LMD) in primary PDACs of treatment-naïve patients^52^. Briefly, a random Forest (RF) classification method was used to build the model setting the number of trees to 500. To avoid overfitting issues, the k-fold cross validation was performed. A selection procedure to maintain only the most informative features was implemented using the recursive feature elimination (rfe) function in caret R package (https://topepo.github.io/caret/). The gene expression and clinical data of the TCGA study cohort were obtained by the GDC database and then the samples were further selected for PDAC diagnosis. Read counts were normalized using DEseq2’s median of ratios. Normalized counts were Log2-transformed and were preprocessed by centering and scaling samples using the Tidymodels R package (https://github.com/tidymodels). PDAC subtypes were predicted in each TCGA sample using the model originated above with the predict function in the randomForest R package setting the parameter type to “class”.

### Differential Expression Analysis

We obtained log-transformed gene-level count data (RSEM expected_count) from the UCSCXenatools^46^, which was subsequently inverse-transformed to raw expected count. TCGA matched normal samples were discarded, and a differential expression (DE) analysis was performed to identify genes with significant expression changes across TCGA-GTEx samples. The DE analysis was conducted using DESeq2^53^, a widely used pipeline for DE analysis. We considered genes with an |LFC|>1 and an adjusted p-value (Padj) < 0.01(Benjamini-Hochberg correction^54^) to be differentially expressed. This process was performed for each of the 16 whole tissue types and 79 subtypes. The contrast for each whole tissue, or subtype, was performed using the corresponding GTEx whole tissue dataset.

### Co-regulation Analysis

To identify co-differentially regulated receptor-ligand pairs, we assigned scores to each of these gene pairs. We considered three major cases: (a) both co-recurrently up/downregulated with significant Padj values (a score of -2/+2), (b) both co-recurrently up/downregulated with at least one of them having a significant Padj value (a score of -1/+1), and (c) no or anti co-differential regulation (a score of 0). This information was displayed in the form of Seaborn clustermap (v0.11.2, Python v3.9.7) with hierarchical clustering performed using the ‘ward’ method of the scipy.cluster library (v1.7.1, Python v3.9.7). On top of this heatmap, G-coupling information (ref Methods section GPCR - G Protein coupling data) and ligand’s mechanism of action (from IUPHAR) was annotated as separate heat maps using matploltib library (v3.4.3, Python v3.9.7) This way we identified differentially expressed genes and co-differentially regulated receptor-ligand pairs in a comprehensive and robust manner. The use of DESeq2 and Benjamini-Hochberg correction ensured the statistical significance of the results, and the consideration of multiple tissue types and subtypes allowed for a more nuanced understanding of the data.

The gene-gene correlations were calculated using the *pearsonr* function from the scipy.stats library in Python v3.9.7. Subsequently, each correlation coefficient was compared to a background distribution of correlation coefficients derived from 1000 randomly selected gene pairs within the respective cancer subtype. The comparison was performed using a t-test from the scipy.stats library in Python v3.9.7. The t-test examines the likelihood that the observed correlation arises from chance variations rather than representing a genuine relationship. A low p-value resulting from the t-test indicates that the correlation coefficient significantly deviates from the expected background distribution. This suggests a potential meaningful association between the genes under investigation.

To account for multiple hypothesis testing, Bonferroni-Hochberg correction was applied to the resulting t-test p-values. The *multipletests* function from the statsmodels library in Python v3.9.7 was utilized for this purpose. A gene pair was considered ’correlated’ if the false discovery rate (FDR) corrected t-test p-value was less than 0.05, and the absolute value of the correlation coefficient exceeded 0.25.

### Gene-Set Enrichment Analysis (GSEA)

We performed gene set enrichment analysis (GSEA) using the WEB-based GEne SeT AnaLysis Toolkit (WebGestalt)^55^. The GSEA process was automated using the R package WebGestaltR. We chose the ’reactome’ pathway database^56^ for enrichment analysis. The results from these analyses were employed to correlate the DE genes to various reactome pathway gene sets with a positive (enriched in TCGA) or negative enrichment (enriched in GTEx) score with Padj<0.01. We considered GSEA results for downstream analyses by filtering pathways containing the GPCR ligands or enzymes belonging to either Signal transduction (‘signaling by GPCR’:R-HSA-372790) or Metabolism (‘metabolism’:R-HSA- 1430728) reactome upper level domains.

### Comparative Transcriptome-Metabolome analysis

We mapped differentially expressed transcripts and metabolites onto a functional interaction network from Reactome pathways^57, 58^, which integrates in the same network information from proteins and small organic molecules. We first collected data from the Pan-Cancer Metabolism Data Explorer^24^, a comprehensive database that contains information about the differential regulation of metabolites associated with cancer aggressiveness. This resource gathered data from over 900 patient samples from five subtypes of cancer: Breast (BRCA), Prostate (PRAD), Pancreas (PAAD), Kidney (KIRC) and Bladder (BLCA). Using this data, we identified the differential expression regulation of enzyme(s) associated with ligand biosynthesis. For this purpose, we used the list of our curated enzymes to query within this external dataset using an HMDB to ChEBI mapping done via MetaboAnalyst^59^. Then, for each pathway, the LFC and Padj values of constituent enzymes were retrieved. Subsequently, this information was merged with the Reactome functional interaction network for a ligand-receptor signaling pathway to complete the entire axis. This was achieved by considering only the first neighbors for involved participants. The resultant networks were used to visualize the directional regulation as well as the DE profiles by utilizing Cytoscape^60^.

### Pathway Abundance Score

We compared the enrichment scores of differentially expressed (DE) transcripts with the differential abundance scores of DE metabolites obtained from the Pan-Cancer Metabolism Data Explorer. To this end, we respectively calculated the GSEA Enrichment Score (ES) and the Pathway Abundance Score (PAS)^24^ by considering only the filtered set of curated biosynthetic pathways from Reactome, specifically for metabolism (R-HSA-1430728). The PAS was calculated by using the formula (I-D)/S, where I represents the number of metabolites (or enzymes) upregulated (LFC>0) in a pathway, D represents the number of metabolites downregulated (LFC<0) in the pathway, and S is the sum of I and D or 1 if I+D=0. A pathway is considered enriched in cancer if the PAS is positive and negative otherwise (similar to ES). We utilized these two metrics to qualitatively evaluate the concordance of enriched biosynthetic pathways in our findings.

### Survival Analysis

The study used univariate unadjusted Cox-Proportional Hazard (Cox-PH) regression models to compute hazard ratios (HR) for predicting the risks of death between high-risk and low-risk TCGA patient groups based on overall survival time. The survival and censoring information was retrieved using TCGAbiolinks for 28 TCGA tissues and 118 subtypes. The risk groups were stratified based on median values of expression, and Kaplan-Meier (KM) plots were used to compare their survival curves. ’survival’ (v4.3.0) and ’survminer’ packages (v0.4.9) in R (v4.2) were used for survival analyses, and statistical significance between survival curves was estimated using log-rank tests with p-values less than 0.05 considered significant.

For the individual components (i.e., Receptors/Ligands/Enzymes), patients were stratified in two risk groups based on the median expression value of each component and HR were estimated. Using the ‘low expression’ group as reference for HR calculations, we label the resulting risk groups as ‘High Survival’ (HR>1) or ‘Low Survival’ (0<HR<1). We further employed the combined expression levels for GPCR axes components i.e., GPCR-ligand/GPCR-enzyme for risk stratification. The patients were now stratified in the two risk groups utilizing the median cutoffs of both the components for e.g., for a GPCR-Ligand based stratification, patients with expression value, EV>MedianGPCR and EV>MedianLigand were placed in one group and the patients with EV<MedianGPCR and EV<MedianLigand in another group. We then filtered out those significant GPCR-axis pairs wherein the Hazard-Ratio axis (HR_axis_) was more than two-fold of the HR in both the individual stratifications (i.e., for above example [HRaxis/HRGPCR<0.5 or HRaxis/HRGPCR>2] and [HRaxis/HRLigand<0.5 or HRaxis/HRLigand>2]) along with the criteria that the logrank-p_axis_ is smaller than the logrank-p values value of both of the individual stratifications and significant i.e., logrank-paxis<0.05. These instances were then displayed on a heatmap created using Seaborn scatterplot created using matplotlib (v3.4.3, Python v3.9.7) with hierarchical clustering performed using the ‘ward’ method of the scipy.cluster library (v1.7.1, Python v3.9.7).

Survival analyses were performed considering whole cancer tissues, as defined in Xena browser, or at the TCGA molecular subtype level.

### Analysis of GPCR drugs on PRISM drug repurposing resource

We retrieved GPCR ligands with cancer growth-inhibitory capacity from a publicly available resource, i.e., PRISM (Program for Response to Inter- and Intra-Species Variation in Cancer Pharmacology)^28^. This drug repurposing resource was developed to identify potential drugs for cancer treatment. To predict drug response, PRISM utilizes genomic data from cancer cell lines and patient-derived xenografts, which are inputted into machine learning pipelines. By taking into account inter- and intra-species variability in drug response, PRISM aims to provide personalized treatment options for specific cancer subtypes. The dataset retrieved from PRISM contained information on the growth-inhibitory activity of over 4,500 drugs tested on 578 human cancer cell lines. This data was filtered based on the drugs which had GPCRs as targets and a list of 52 GPCR drugs/ligands was obtained. To identify active drugs for cancer treatment, PRISM uses the bimodality coefficient and correlation as metrics. The bimodality coefficient measures the separation between the two peaks in a drug response distribution, identifying drugs with a bimodal distribution of response across cancer cell lines. The correlation measures the strength of the relationship between predicted and actual drug response in patient-derived xenograft models. Following a similar approach, we used these metrics (correlation>0.2, bimodality coefficient>0.35) to identify GPCR drugs that are active against specific cancer subtypes. The active drugs were then investigated on the basis of GPCRs associated with survival in either of GPCR-Ligand or GPCR-Enzyme axes. The information was further annotated with G-coupling information and ultimately displayed using visual representations such as scatter plots, venn diagram and heatmaps utilizing seaborn (v0.11.2, Python v3.9.7) and matplotlib libraries (v3.4.3, Python v3.9.7).

We also considered data obtained from two recent PRISM Repurposing screens: Repurposing-1M and Repurposing-300. These screens assess the Log Fold Change (LFC) in cell counts, deploying the PRISM assay to evaluate 6658 compounds, of which 1514 target GPCRs. The compounds are administered at a dose of 2.5 μM over a 5-day treatment to 906 cancer cell lines. The final LFC values are consolidated in the https://depmap.org/portal/download/all/?release=PRISM+Repurposing+Public+23Q2&file=R epurposing_Public_23Q2_Extended_Primary_Data_Matrix.csv table, constituting a matrix where rows represent individual treatments and columns correspond to the DepMap IDs. For convenience, this matrix also includes the compounds from the original PRISM Repurposing Primary screen. The research also incorporates metadata for the drugs and the cell lines from Repurposing_Public_23Q2_Extended_Primary_Data_Matrix.csv and the Model.csv file, respectively. Using the Model.csv, we leverage DepMapID to stratify cell lines according to tissue type, following the OncoTree ontology. All of this data is freely accessible on the DepMap portal. For each cell line, we considered the top 10 GPCR drugs inhibiting cancer cell growth, i.e. having the most negative log2fold change. We then performed statistic considering the Mechanism of Action (MOA) and Targets associated to each ligand.

## Data Availability

Code and data used for this study are available at: https://gpcrcanceraxes.bioinfolab.sns.it/

## Conflict of interest

JSG reports consulting fees from Domain Pharmaceuticals, Pangea Therapeutics, and io9, and is founder of Kadima Pharmaceuticals, all unrelated to the current study.

## Acknowledgments

F.R. was supported by the Italian Ministry of University and Research through the Department of excellence “Faculty of Sciences” of Scuola Normale Superiore. The research leading to these results also received funding from the Italian Association for Cancer Research (AIRC) under My First AIRC Grant (MFAG) 2020 - ID. 24317 project – P.I. F.R; the US National Institutes of Health (U24 HG012198) to G.W. We gratefully acknowledge the CINECA award, in collaboration with AIRC, for the availability of high performance computing resources and support. We gratefully acknowledge computational resources of the Center for High Performance Computing (CHPC) at Scuola Normale Superiore.

## Supplementary Figure s

**Supplementary** Figure 1: a) stacked barplot with the distribution of DE GPCRs for each cancer tissue categorized by coupling information; b) heatmap showing the LFCs from DE analysis (TCGA vs GTEX, via DESEQ2) of G protein in each cancer tissue. Significant instances (BH Padj<0.01) are annotated with “*”; c) same as b) but for cancer molecular subtypes; d) heatmap showing ligand-associated pathways which are significantly enriched in at least one cancer subtype. An enriched pathway refers to a pathway enriched in TCGA subtype with a GSEA enrichment score>0 and BH corrected Padj<0.01 .

**Supplementary** Figure 2: Heatmap displaying the co-differential regulation for Receptor-Ligand pairs across different TCGA subtypes (color-coded at the top row). Darker red represents both receptor and ligand significantly co-up regulated in TCGA (i.e., LFC>1 and BH corrected Padj<0.01) and darker blue represents both receptor and ligand significantly co-down regulated in TCGA (i.e., LFC<1 and BH corrected Padj<0.01). Paler colors represent either of the receptor-ligand as significantly DE (i.e., |LFC|>1 for both but BH corrected Padj<0.01 for only one of these). White cells indicate anti-regulation or no significant fold change (|LFC|<1) at all in at least one of them.

**Supplementary** Figure 3: functional interaction network between genes (ovals) and metabolites (diamonds) in the ‘Purine catabolism’ pathway in a) pancreatic and b) prostatic cancer. Red nodes indicate upregulated components in cancer, blue nodes indicate downregulated components, and green nodes indicate no information available. The network shows that over-activation of the pathway is contributed by over-expression of both genes and metabolites.

**Supplementary** Figure 4: Heatmap (center panel) displaying the co-differential regulation for Receptor-Enzyme pairs across different TCGA subtypes (color-coded at the top row). Darker red represents both receptor and ligand significantly co-up regulated in TCGA (i.e., LFC>1 and Padj<0.01) and darker blue represents both receptor and ligand significantly co-down regulated in TCGA (i.e., LFC<1 and Padj<0.01). Paler colors represent either of the receptor-ligand as significantly DE (i.e., |LFC|>1 for both but Padj<0.01 for only one of these). White cells indicate anti-regulation or no fold change at all in at least one of them. Only those pairs which are affected in at least 25% of TCGA subtypes are displayed. A dashed line separating the two clusters, created using Hierarchical clustering, is shown in the middle. Hierarchical clustering was performed using the ’ward’ method to identify two homogeneous clusters by minimizing within-cluster variance based on the sum of squared differences of feature values i.e. scores∈[-2,+2] assigned to each pair. Heatmap (left panel) uses color codes to display the ligand’s mechanism of action in ‘Action’, G Protein-coupling associated with the GPCRs and GPCR Family.

**Supplementary** Figure 5: barplot with the TCGA molecular subtype-level statistics of the receptor-enzyme pairs with significant receptor-enzyme correlation.

**Supplementary** Figure 6:Kaplan-Meier curve for survival analysis done by stratifying patients based on expression values of representative GPCR-ligand axes, found to be more significantly associated with lower survival than individual components in several cancer types. Each expression group is represented with a colored survival curve, as denoted in the legend. Risk tables below the KM plots represent the number of surviving patients at various time-points. Logrank-p values are annotated within the KM plots.

**Supplementary** Figure 7: Kaplan-Meier curve for survival analysis done by stratifying patients based on expression values of SSTR2-SST axis in either head and neck or adrenal gland cancers. Each expression group is represented with a colored survival curve, as denoted in the legend. Risk tables below the KM plots represent the number of surviving patients at various time-points. Logrank-p values are annotated within the KM plots.

**Supplementary** Figure 8:G protein coupling preferences of axes found significantly associated with higher survival in head and neck: i.e., *SSTR2-SST*, *RXFP3-INSL5* and *RXFP1-RLN3*

**Supplementary** Figure 9: The bubble plot shows the correlation between the combined-expression levels of GPCR-ligand pairs and patient survival across cancer subtypes. The pairs with a log rank p-value <0.05 and also lower than log rank p-values for individual GPCR/ligand are displayed. Bubble color is proportional to HR: i.e., HR>1 : High expression is correlated with high survival (red); HR<1: High expression is correlated with poor survival (blue). Bubble diameters are proportional to the -log10(log-rank p-value). Green highlighted bubbles represent the most significant instances (sample sizes>5, FDR<0.1).

**Supplementary** Figure 10: The bubble plot shows the correlation between the combined-expression levels of GPCR-enzyme pairs and patient survival across various subtypes. The pairs with a log rank p-value <0.05 and also lower than log rank p-values for individual GPCR/ligand are displayed. Bubble color is proportional to HR: i.e., HR>1 : High expression is correlated with high survival (red); HR<1: High expression is correlated with poor survival (blue). Bubble diameters are proportional to the -log10(log-rank p-value). Green highlighted bubbles represent the most significant instances (sample sizes>5, FDR<0.1).

**Supplementary** Figure 11: frequency of Pubmed evidences obtained by querying with the following keywords: a) GPCR name *and* ligand name *and* cancer;b) GPCR name *and* ligand name *and* cancer *and* tissue; c) GPCR name *and* ligand name *and* cancer *and* immunotherapy; d) GPCR name *and* enzyme name *and* cancer

## Supplementary Tables

**Supplementary Table 1**: GPCR ligands in Reactome Pathways (Metabolism and Signal Transduction domains)

**Supplementary Table 2**: List of GPCR ligands with cognate receptors and biosynthetic enzymes from RHEADB

**Supplementary Table 3**: Tissue-wise sample distribution in the data retrieved from UCSCXena.

**Supplementary Table 4**: Reactome pathways enrichment scores (ES) and differential abundance scores (PAS) calculated from transcripts and metabolites log2fold changes

**Supplementary Table 5**: Expression correlation statistics for receptor-enzyme pairs

**Supplementary Table 6**: GPCR based survival analysis - cancer tissue/TCGA molecular subtypes

**Supplementary Table 7**: ligand based survival analysis - cancer tissue/TCGA molecular subtypes

**Supplementary Table 8**: enzyme based survival analysis - cancer tissues/TCGA molecular subtypes

**Supplementary Table 9**: GPCR-ligand based survival analysis - cancer tissues/TCGA molecular subtypes

**Supplementary Table 10**: GPCR-enzyme based survival analysis - cancer tissues/TCGA molecular subtypes

**Supplementary Table 11**: GPCR ligands activity from PRISM cancer cell line drug repurposing hub (https://depmap.org/repurposing/)

**Supplementary Table 12**: GPCR ligands activity (log2Fold Change) from PRISM Repurposing Public 23Q2

**Supplementary Table 13**: Literature evidence for GPCR axes

**Supplementary Table 14**: Literature support for rate-limiting role of Metabolism pathways’s enzymes

## Notes

https://gpcrcanceraxes.bioinfolab.sns.it/

### Competing Interest Statement

The authors have declared no competing interest.

### Summary of Updates

Updated analysis of the PRISM dataset included

